# Domestication effects on aggressiveness: Comparison of biting motivation and bite force between wild and domesticated finches

**DOI:** 10.1101/2021.02.24.432800

**Authors:** Kenta Suzuki, Kazuo Okanoya

## Abstract

Domesticated animals evolve unique traits, known as domestication phenotypes or the domestication syndrome, due to their adaptation to a captive environment and changes in selection pressures. After being tamed, the Bengalese finch (*Lonchura striata* var. *domestica*) has undergone behavioural and physiological trait changes that differ from those of its wild ancestor, the white-rumped munia (*Lonchura striata*). The Bengalese finch has complex songs, lower fear response, and lower corticosterone levels than those in the white-rumped munia. We hypothesized that domesticated finches increase the effort to reproduce in lieu of maintaining fitness for survival as needed in the wild. Aggressiveness and bite performance affect survival rates and reproductive success, and are good indicators of adaptability in the natural environment. Therefore, we compared the aggressiveness and biting force of white-rumped munias with those of Bengalese finches to explore the evolutionary mechanisms of behavioural changes due to domestication. We found that the Bengalese finch had decreased aggressiveness (incidence of aggressive biting birds and the number of bite responses) and bite force than those in the white-rumped munia due to domestication. Therefore, we believe they could allocate more resources for breeding that would otherwise be needed for coping with predators through aggression.

## 1. Introduction

The domestication process involves adaptation to a captive environment and a shift in selection pressures (Price, 1984, 1999). Such domestication processes, now called domestication phenotypes or the domestication syndrome, alter the morphology, physiological status, and behavioural characteristics of domesticated animals (Darwin, 1868; Hale, 1969; Price, 1984; Künzl and Sachser, 1999; Lepage et al., 2000; Wilkins et al., 2014). However, most of these traits have been reported anecdotally without any accompanying frequencies or measurements (Load et al., 2020). Animal domestication is useful for studying the evolutionary mechanisms of traits in response to changes in selection pressure. Understanding the process requires an approach focused on essential adaptations to artificial environments where, although specific adaptations may vary between species, the selective pressures are common for all (Load et al., 2020).

The Bengalese finch (*Lonchura striata* var. *domestica*) has been domesticated from its wild ancestor, the white-rumped munia (*Lonchura striata*) that was originally found in China (Washio, 1996; Okanoya, 2004a, b). There are behavioural differences in the song complexity and fear response of the Bengalese finch and the white-rumped munia, and physiological differences in their stress hormone corticosterone levels. The Bengalese finch has phonologically and syntactically complex songs with louder and pure tone-like sounds (Honda and Okanoya, 1999), lower fear response (Suzuki et al., 2013), and lower corticosterone levels (Suzuki et al., 2012, 2014a, b) compared to those in the white-rumped munia. These variations of the traits in finches are useful model systems for analysing the evolutionary mechanisms of domestication.

Aggressiveness is a trait that affects fitness in animals/birds and has been associated with survival and reproductive success. More aggressive individuals are better able to survive and reproduce in the wild (Biro and Stamps, 2008; Ariyomo and Watt, 2012). Aggressive behaviour is used to defend a resource and territory, defend one’s self against predators, and compete for mates. In many animals, defensive attack, freezing, and fleeing are common defensive behaviours against predators (Eilam, 2005). Maintaining high aggression is important because attacking predators and rivals is the last resort for an animal to protect itself, its mate, and their offspring. However, investment in such traits is often costly as the expression of such traits requires energy and reduces self-maintenance and the production and care of offspring (Cain and Ketterson, 2013).

Furthermore, bite performance (bite force) is a useful trait related to an individual’s fitness for survival in the wild, such as for food acquisition, defence against predators, and aggressive interactions (Anderson et al., 2008; Vicenzi et al., 2020). Bite force is correlated to body size and beak depth in finches (Van der Meij and Bout, 2004; Herrel et al., 2005). Strongly biting individuals are considered more adaptive in nature because they can crack harder seeds (Herrel et al., 2005). Therefore, aggressiveness and bite force can be used as good indicators of adaptability in the natural environment.

We hypothesized that domesticated birds exhibited changed behaviours related to their fitness for survival, such as defensive behaviour against predators, by adaptation to artificial, non-predatory, and food-rich environments, and that they increased their efforts to reproduce (e.g. complex song development) in exchange for maintaining their fitness for survival in the wild. In the present study, we compared the aggressiveness and biting force of wild finches with those of domesticated finches to explore the evolutionary mechanisms of behavioural changes that occur due to domestication.

## 2. Method

### 2.1. Subjects and husbandry

We used 63 adult Bengalese finches (32 males and 31 females) and 37 adult white-rumped munias (19 males and 18 females). The Bengalese finches were bred in our laboratory. The white-rumped munias were bought from a commercial supplier (n = 13), captured in the wild in Taiwan (n = 2), or bred in our laboratory (n = 22). These birds were housed in six groups (each species and sex) in the same type of cages (cage size: 370 × 415 × 440 mm) in a animal room at RIKEN Brain Science Institute, Tokyo, Japan under a 13 h light and 11 h dark cycle, 25 °C ambient temperature, and 50 % humidity. Seed mixture, shell grit, and vitamin-enhanced water were available *ad libitum* to the birds. All experimental procedures and the housing conditions of the birds were approved by the Animal Experiments Committee at RIKEN (#H24-2-229), and all the birds were cared for in accordance with the Institutional Guidelines of RIKEN for Experiments Using Animals.

### 2.2. The measurement of biting motivation and performance

We measured biting motivation and performance in Bengalese finches and white-rumped munias using a Tekscan FlexiForce ELF™ system (Economical Load & Force Measurement System; Tekscan Inc., MA, USA) with a piezoresistive force sensor (model #B201; a strip of thin plastic 10 mm wide, 150 mm long, and 0.2 mm thick). This system has previously been used to measure bite force in small animals such as insects and mammals (Freeman and Lemen, 2008; Hall et al., 2010). Before the test, the sensor was calibrated by applying known forces using a digital force gauge (model FGP-2; Nidec-Shimpo Corp., Japan). The measuring range was between 0 and 10.82 N. The birds were caught from the rearing cage one by one and immediately subjected to the test. We held a bird in the hand and moved its beak close to the sensor placed on the desk and observed for 5 seconds. We measured the number of biting responses and the bite force. Five continuous measurements (the bird was brought near the sensor five times) were repeated for each individual. The average values of bite force (mean bite force) and maximal values of bite force (maximal bite force) from five bite trials were used for analysis. If the bird did not show biting behaviour, the bite force of that trial was not used for analysis. At the end of the test, we measured the body weight of the birds to the nearest 0.1 g.

### 2.3. Statistical analysis

Differences between strains in the incidence of birds biting the sensor stick were tested using a Chi-square (χ^2^) test. The number of bite responses was analysed using the Mann-Whitney *U* test with Bonferroni correction, as count variables that did not approximate a normal distribution. The mean and maximum bite forces were analysed by two-way ANOVA (factors: strain and sex). A significance level of *P* < 0.05 was used for all statistical analyses. Statistical analyses were performed using the Stat View software ver. 5.0 (SAS Institute Inc., NC, USA).

## 3. Results

The incidence of aggressive biting was significantly higher in white-rumped munias (32/37 birds, 86.5 %) than that in Bengalese finches (33/63 birds, 52.4 %) (OR: 5.82, 95 % CI: 2.06 - 16.28, χ^2^ = 11.9, *P* < 0.001). The number of bite responses was significantly affected by strain (*Z* = −5.21, Bonferroni adjusted *P* < 0.001), but there was no effect of sex (*Z* = −1.82, Bonferroni adjusted *P* = 0.27) and strain × sex interactions (Bengalese finch: *Z* = −2.43, Bonferroni adjusted *P* = 0.09; white-rumped munia: *Z* = − 0.66, Bonferroni adjusted *P* > 0.99). The biting responses (number of bites per five trials) were lower in Bengalese finches [mean = 1.51, SE = 0.23, median (25-75 percentiles) = 1.00 (0.00-2.00)] than those in white-rumped munias [mean = 4.14, SE = 0.30, median (25-75 percentiles) = 5.00 (5.00-5.00)] (Fig. 1).

**Fig. 1.**
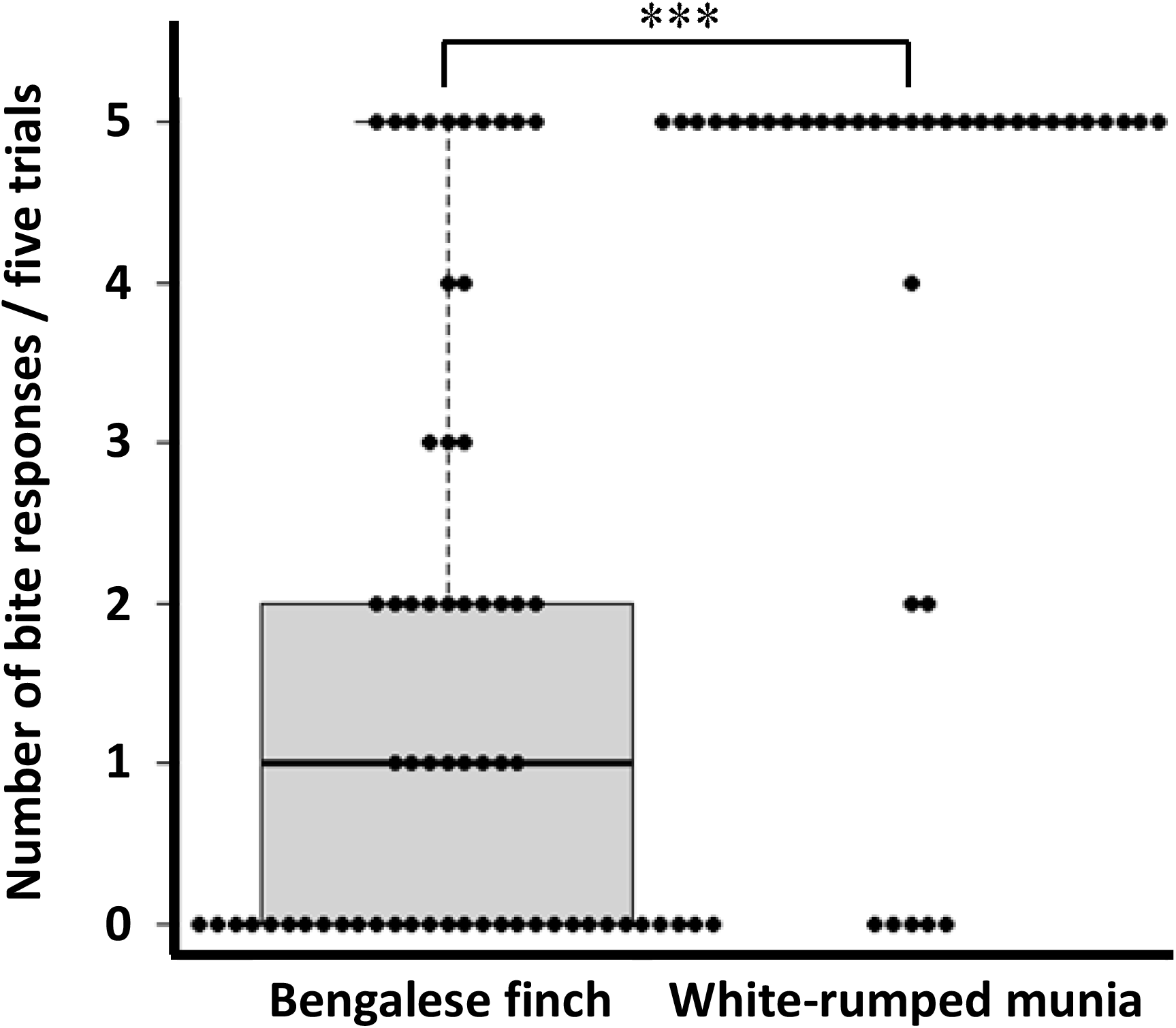
Number of biting responses per five trials in Bengalese finches (*Lonchura striata* var. *domestica*) and white-rumped munias (*Lonchura striata*). The biting responses were significantly lower in Bengalese finches [median (25-75 percentiles), 1.00 (0.00-2.00)] than those in white-rumped munias [median (25-75 percentiles), 5.00 (5.00-5.00)], \*\*\**P* < 0.001.

The mean bite force (average values of all trials) showed the effect of strain (*F* = 21.8, *P* < 0.0001), but not sex (*F* = 0.01, *P* = 0.92) or strain × sex interactions (*F* = 3.71, *P* = 0.06). The mean bite force was stronger in white-rumped munias (mean = 2.03 N, SE = 0.12) than that in Bengalese finches (mean = 1.20 N, SE = 0.14) (Fig. 2A). The maximal bite force (maximum values of all trials) showed the effect of strain (*F* = 37.2, *P* < 0.0001) but not that of sex (*F* = 0.04, *P* = 0.84). The interactions of strain × sex showed a significant effect on bite force (*F* = 5.11, *P* = 0.03) but the Bonferroni/Dunn post-hoc test showed no significant difference (*P* > 0.05). Maximal bite force was stronger in white-rumped munias (mean = 3.26 N, SE = 0.20) than that in Bengalese finches (mean = 1.57 N, SE = 0.21) (Fig. 2B).

**Fig. 2.**
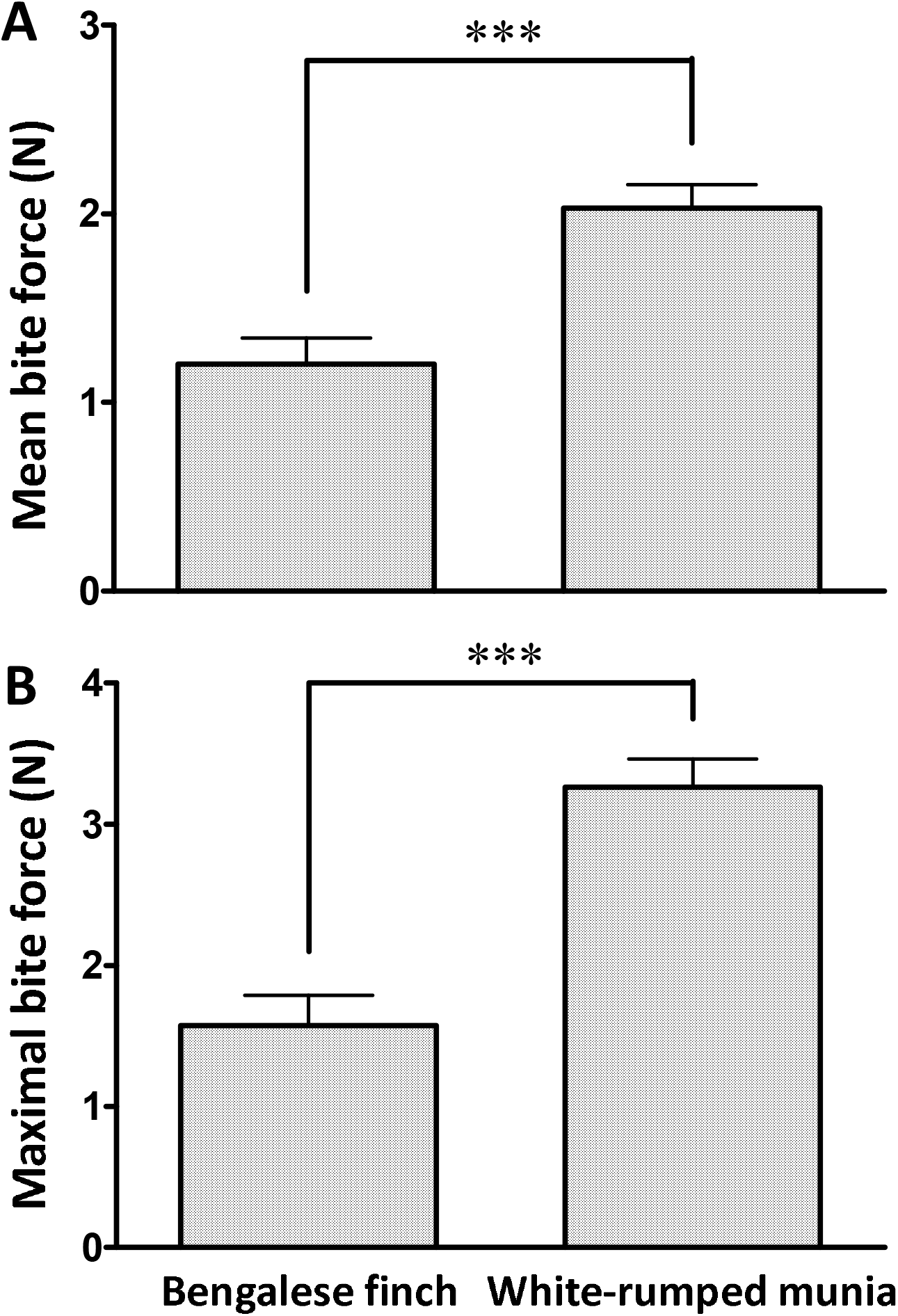
(A) Mean and (B) maximal values of bite force in Bengalese finches (*Lonchura striata* var. *domestica*) and white-rumped munias (*Lonchura striata*). Bars are means and vertical lines are standard errors. Bengalese finches have significantly lower mean and maximal bite forces than those in white-rumped munias (\*\*\**P* < 0.001).

Body weights indicated a significant effect of strain by two-way ANOVA (*F* = 56.94, *P* < 0.0001). The Bengalese finches had a greater mean body weight than that of the white-rumped munias (Bengalese finch: mean = 14.25 g, SE = 0.21; white-rumped munia: mean = 11.27 g, SE = 0.19). We found a significant interaction effect of strain × sex (F = 4.447, *P* < 0.05) on body weight. Female Bengalese finches were heavier than male birds (male: mean = 13.72 g, SE = 0.28; Female: mean =14.78 g, SE = 0.29; Bonferroni/Dunn test, *P* < 0.05). However, no differences in body weight based on sex were found in white-rumped munias (Male: mean = 11.48 g, SE = 0.32; Female: mean = 11.03 g, SE = 0.19; Bonferroni/Dunn test, *P* > 0.05).

## 4. Discussion

We examined bite response and biting force as a measure of aggressiveness to explore the mechanisms of behavioural changes due to domestication in the Bengalese finch. Our results show that domesticated Bengalese finches’ aggressiveness was lower than that of their wild ancestor, white-rumped munias. The Bengalese finches showed a lower incidence of aggressive biting birds, number of bite responses per five trials (Fig. 1), and bite force (Fig. 2) compared to those in white-rumped munias. The bite response and bite force showed no relationship with sex in either strain. Therefore, we determined that Bengalese finches exhibit decreased aggressiveness due to domestication.

Maximal bite force has been found to be repeatable among individuals across all vertebrates studied (Anderson et al., 2008). Therefore, it is an effective indicator of individual traits. Furthermore, maximal bite force is strongly correlated with body and head sizes in many taxa (Anderson et al., 2008). Van der Meji and Bout (2004) reported that maximal bite force correlates positively with body mass in finches. Species related closely to white-rumped munia showed similar maximal bite forces [Pale-headed munia (*Lonchura pallida*): 3.3 N, Scaly breasted munia (*Lonchura punctulata*): 3.7 N {Van der Meji and Bout, 2004}; white-rumped munia: 3.26 N {our study}], because these species have similar body mass [Pale-headed munia: 13.2 g, Scaly breasted munia: 12.2 g (Van der Meji and Bout, 2004); white-rumped munia: 11.3 g (our study)]. However, in our findings, the Bengalese finches showed lower maximal bite force (1.57 N) than those in white-rumped munias despite heavier body mass (Bengalese finch: 14.3 g). These results show that the Bengalese finches have lower biting force due to domestication, regardless of their higher body weight.

Domesticated animals commonly exhibit decreased levels of aggression, including intraspecific, interspecific, offensive, or defensive aggression (Künzl and Sachser, 1999; Hare et al., 2012). Bengalese finches were artificially selected for their reproductive efficiency, parental behaviour, and ability to foster the offspring of other bird species, and their white colour morphs (Taka-Tsukasa, 1917). In the process of domestication, nervous individuals that affect reproductive efficiency may have been eliminated and individuals with low fear of humans may have been selected. Bengalese finches exhibited a decreased fear response (tonic immobility response) compared with that in white-rumped munias (Suzuki et al., 2013). Therefore, it is considered that the Bengalese finch was not afraid of humans and therefore expressed less aggression, even in the experimental situation of this study. In another study, the experimental selection of red junglefowls with reduced fear of humans resulted in expression of the typical domestication phenotypes acquired in a few generations (Agnvall et al., 2018; Katajamaa and Jensen, 2020). Therefore, tameness with a reduction in avoidance responses to humans may have driven the evolution of domesticated phenotypes.

Some reports indicate that male aggression and paternal care are associated, and that aggressive males provide poor paternal care (Qvarnstr◻m, 1997). Male aggression of Western bluebird (*Sialia mexicana*) was negatively related to reproductive success through its effects on male parental care (Duckworth, 2006). Aggressive Gouldian finch (*Chloebia gouldiae*) males provided parental effort comparable to that of non-aggressive males in a low competition environment, but severely reduced or abandoned parental investment in highly competitive environments (Pryke and Griffith, 2009). Therefore, aggressive traits are considered costly in a natural environment. The relationship between aggressiveness and reproductive efficiency is likely a trade-off. Bengalese finches may have improved reproductive efficiency, such as increasing song complexity and paternal care, by reducing the cost of coping with predators due to domestication.

In conclusion, we found that domesticated Bengalese finches show decreased aggressiveness compared with that in their wild ancestor, the white-rumped munia. We believe that they can allocate more resources for reproduction (good breeding and parenting) in exchange for the ability to cope with predators (aggressiveness and fearfulness). Further studies are needed to understand the proximate mechanisms underlying these trade-offs. The findings of this study may help elucidate the biological basis for the evolution of language and reduced aggression due to human self-domestication.

## Acknowledgements

We thank Dr. Yuko Ikkatai and Ms. Keiko Asai (RIKEN Brain Science Institute, Tokyo, Japan) for animal care. We are grateful to Dr. Ryosuke O. Tachibana (The University of Tokyo, Tokyo, Japan) and Dr. Hiroko Kagawa (JST-ERATO Okanoya Emotional Information Project, Tokyo, Japan) for their technical advice. We thank Ms. Tomoko Mizuhara (The University of Tokyo, Tokyo, Japan) for experimental assistance. This study was supported by JST-ERATO and RIKEN Brain Science Institute.

## Author contributions

**Kenta Suzuki:** Conceptualization, Data curation, Formal analysis, Investigation, Methodology, Visualization, Writing - Original draft; Kazuo Okanoya: Conceptualization, Funding acquisition, Supervision.

## References

Agnvall, B., Bélteky, J., Katajamaa, R., Jensen, P., 2018. Is evolution of domestication driven by tameness? A selective review with focus on chickens. Appl. Anim. Behav. Sci. 205, 227–233. https://doi.org/10.1016/j.applanim.2017.09.006.

Anderson, R. A., McBrayer, L. D., Herrel, A., 2008. Bite force in vertebrates: opportunities and caveats for use of a nonpareil whole-animal performance measure. Biol. J. Linn. Soc. 93(4), 709–720. https://doi.org/10.1111/j.1095-8312.2007.00905.x.

Ariyomo, T. O., Watt, P. J., 2012. The effect of variation in boldness and aggressiveness on the reproductive success of zebrafish. Anim. Behav. 83(1), 41–46. https://doi.org/10.1016/j.anbehav.2011.10.004.

Biro, P. A., Stamps, J. A., 2008. Are animal personality traits linked to life-history productivity?. Trends Ecol Evol., 23(7), 361–368. https://doi.org/10.1016/j.tree.2008.04.003.

Cain, K. E., Ketterson, E. D., 2013. Costs and benefits of competitive traits in females: aggression, maternal care and reproductive success. PLoS One, 8(10), e77816. https://doi.org/10.1371/journal.pone.0077816.

Darwin, C., 1868. The variation of animals and plants under domestication. John Murray, London.

Duckworth, R. A., 2006. Behavioral correlations across breeding contexts provide a mechanism for a cost of aggression. Behav. Ecol. 17(6), 1011–1019. https://doi.org/10.1093/beheco/arl035.

Eilam, D., 2005. Die hard: a blend of freezing and fleeing as a dynamic defense—implications for the control of defensive behavior. Neurosci. Biobehav. Rev. 29(8), 1181–1191. https://doi.org/10.1016/j.neubiorev.2005.03.027.

Freeman, P. W., Lemen, C. A., 2008. Measuring bite force in small mammals with a piezo-resistive sensor. J. Mammal. 89(2), 513–517. https://doi.org/10.1644/07-MAMM-A-101R.1.

Hale, E.B., 1969. Domestication and the evolution of behaviour. in: Hafez, E.S.E. (Ed) The Behaviour of Domestic Animals, 2nd Edition. Baillière, Tindall and Cassell, London, pp. 22–42.

Hall, M. D., McLaren, L., Brooks, R. C., Lailvaux, S. P., 2010. Interactions among performance capacities predict male combat outcomes in the field cricket. Funct. Ecol., 24(1), 159–164. https://doi.org/10.1111/j.1365-2435.2009.01611.x.

Hare, B., Wobber, V., Wrangham, R., 2012. The self-domestication hypothesis: evolution of bonobo psychology is due to selection against aggression. Anim. Behav. 83(3), 573–585. https://doi.org/10.1016/j.anbehav.2011.12.007.

Herrel, A., Podos, J., Huber, S. K., Hendry, A. P., 2005. Bite performance and morphology in a population of Darwin’s finches: implications for the evolution of beak shape. Funct. Ecol., 19(1), 43–48. https://doi.org/10.1111/j.0269-8463.2005.00923.x.

Honda, E., Okanoya, K., 1999. Acoustical and syntactical comparisons between songs of the white-backed munia (*Lonchura striata*) and its domesticated strain, the Bengalese finch (*Lonchura striata* var. *domestica*). Zool. Sci. 16, 319–326. https://doi.org/10.2108/zsj.16.319.

Katajamaa, R., Jensen, P., 2020. Tameness correlates with domestication related traits in a Red Junglefowl intercross. Genes Brain Behav. e12704. https://doi.org/10.1111/gbb.12704.

Künzl, C., Sachser, N., 1999. The behavioral endocrinology of domestication: A comparison between the domestic guinea pig (*Cavia apereaf.porcellus*) and its wild ancestor, the cavy (*Cavia aperea*). Horm. Behav. 35, 28–37. https://doi.org/10.1006/hbeh.1998.1493.

Lepage, O., Øverli, Ø., Petersson, E., Järvi, T., Winberg, S., 2000. Differential stress coping in wild and domesticated sea trout. Brain Behav. Evol. 56, 259–268. https://doi.org/10.1159/000047209.

Lord, K. A., Larson, G., Coppinger, R. P., Karlsson, E. K., 2020. The history of farm foxes undermines the animal domestication syndrome. Trends Ecol. Evol. 35(2), 125–136. https://doi.org/10.1016/j.tree.2019.10.011.

Okanoya, K., 2004a. Song syntax in Bengalese finches: proximate and ultimate analyses. Adv. Stud. Behav. 34, 297–346. https://doi.org/10.1016/S0065-3454(04)34008-8.

Okanoya, K., 2004b. The Bengalese finch: a window on the behavioral neurobiology of birdsong syntax. Ann. N. Y. Acad. Sci. 1016, 724–735. DOI:10.1196/annals.1298.026.

Price, E.O., 1984. Behavioral aspects of animal domestication. Q. Rev. Biol. 59, 1–32. https://www.jstor.org/stable/2827868.

Price, E. O., 1999. Behavioral development in animals undergoing domestication. Appl. Anim. Behav. Sci. 65(3), 245–271. https://doi.org/10.1016/S0168-1591(99)00087-8.

Pryke, S. R., Griffith, S. C., 2009. Socially mediated trade-offs between aggression and parental effort in competing color morphs. Am. Nat. 174(4), 455–464. https://doi.org/10.1086/605376.

Qvarnström, A., 1997. Experimentally increased badge size increases male competition and reduces male parental care in the collared flycatcher. Proc. Royal Soc. B 264(1385), 1225–1231. https://doi.org/10.1098/rspb.1997.0169.

Suzuki, K., Ikebuchi, M., Bischof, H. J., Okanoya, K., 2014a. Behavioral and neural trade-offs between song complexity and stress reaction in a wild and a domesticated finch strain. Neurosci. Biobehav. Rev. 46, 547–556. https://doi.org/10.1016/j.neubiorev.2014.07.011.

Suzuki, K., Ikebuchi, M., Okanoya, K., 2013. The impact of domestication on fearfulness: a comparison of tonic immobility reactions in wild and domesticated finches. Behav. Proc. 100, 58–63. https://doi.org/10.1016/j.beproc.2013.08.004.

Suzuki, K., Matsunaga, E., Yamada, H., Kobayashi, T., Okanoya, K., 2014b. Complex song development and stress hormone levels in the Bengalese finch. Avian Biol. Res. 7(1), 10–17. https://doi.org/10.3184/175815514X13903270812502.

Suzuki, K., Yamada, H., Kobayashi, T., Okanoya, K., 2012. Decreased fecal corticosterone levels due to domestication: a comparison between the white-backed munia (*Lonchura striata*) and its domesticated strain, the Bengalese finch (*Lonchura striata* var. *domestica*) with a suggestion for complex song evolution. J. Exp. Zool. A 317, 561–570. https://doi.org/10.1002/jez.1748.

Taka-Tsukasa, N., 1917. Companion birds. Shoka-bo, Tokyo. (in Japanese)

Van der Meij, M. A. A., Bout, R. G., 2004. Scaling of jaw muscle size and maximal bite force in finches. J. Exp. Biol., 207(16), 2745–2753. https://doi.org/10.1242/jeb.01091.

Vicenzi, N., Laspiur, A., Sassi, P. L., Massarelli, R., Krenz, J., Ibargüengoytía, N. R., 2020. Impact of temperature on bite force and bite endurance in the Leopard Iguana (*Diplolaemus leopardinus*) in the Andes Mountains. J. Exp. Biol. 223, jeb221382. https://doi.org/10.1242/jeb.221382.

Washio, K., 1996. Chasing the mystery of the Bengalese finch. Modern Literature Inc., Tokyo. (in Japanese)

Wilkins, A. S., Wrangham, R. W., Fitch, W. T., 2014. The “domestication syndrome” in mammals: a unified explanation based on neural crest cell behavior and genetics. Genetics, 197(3), 795–808. https://doi.org/10.1534/genetics.114.165423.

